# MFGE8 links absorption of dietary fatty acids with catabolism of enterocyte lipid stores through HNF4γ-dependent transcription of CES enzymes

**DOI:** 10.1101/2022.05.16.492147

**Authors:** Ritwik Datta, Mohammad A Gholampour, Christopher D Yang, Regan Volk, Sinan Lin, Michael J Podolsky, Thomas Arnold, Florian Rieder, Balyn W. Zaro, Michael Verzi, Richard Lehner, Nada Abumrad, Carlos O Lizama, Kamran Atabai

## Abstract

Enterocytes modulate the extent of postprandial lipemia, a potent risk factor for developing atherosclerotic disease, by storing dietary fats in cytoplasmic lipid droplets (cLDs). We have previously demonstrated that the integrin ligand MFGE8 links absorption of dietary fats with activation of triglyceride (TG) hydrolases that catabolize cLDs for chylomicron production. The hydrolase(s) responsible for mobilization of TG from diet-derived cLDs is unknown though recent evidence indicates that this process is independent of the canonical pathway of TG hydrolysis mediated by ATGL. Here we identify CES1D as the key hydrolase downstream of the MFGE8-αvβ5 integrin pathway that regulates catabolism of diet-drive cLDs. *Mfge8* KO enterocytes have reduced CES1D transcript and protein levels and reduced protein levels of the transcription factor HNF4γ. Mice KO for *Ces1d* or *Hnf4γ* have decreased enterocyte TG hydrolase activity coupled with retention of TG in cLDs. Mechanistically, MFGE8-dependent fatty acid uptake through CD36 leads to stabilization of HNF4γ protein levels; HNF4γ then increases *Ces1d* transcription. Our work identifies a regulatory network by which MFGE8 and αvβ5 regulate the severity of postprandial lipemia by linking dietary fat absorption with protein stabilization of a transcription factor that increases expression of enterocyte TG hydrolases that catabolize diet-derived cLDs.

## INTRODUCTION

Intestinal lipid homeostasis has important implications for the development of atherosclerotic heart disease (1, 2). In addition to absorbing nutrients, the small intestine functions as a lipid storage organ that can limit postprandial serum lipid levels by storing a proportion of absorbed fats in cytoplasmic lipid droplets (cLDs), (3, 4). The clinical relevance of this under-appreciated small intestinal function is evident by the stronger correlation of postprandial lipid levels with coronary artery disease as compared with more commonly measured fasting serum lipid levels (2). In obesity, insulin resistance at the level of the intestine removes the suppressive effect of insulin on chylomicron production resulting in more severe postprandial lipemia (7). Humans with visceral obesity also demonstrate more severe postprandial lipemia and an increased risk of cardiovascular disease (8).

CLDs are increasingly recognized as dynamic organelles with pleotropic functions that include prevention of fatty acid-induced lipotoxicity, serving as platforms for protein binding and degradation, and providing a reservoir for hydrophobic molecules important in numerous cellular functions. The small intestine is unique in that enterocytes contain distinct pools of cLDs derived from dietary fat or from lipids taken up from the basolateral circulation (5, 6). Our current understanding of cytoplasmic lipid droplet (cLD) metabolism is primarily derived from work done in adipocytes and hepatocytes, including the identification of several molecules that associate with and regulate hydrolysis of triglycerides (TG) in cLDs. Adipocyte triglyceride lipase (ATGL) is the predominant intracellular hydrolase responsible for cleaving intracellular TG to diacylglycerol (7) and *Atgl* KO mice accumulating TG in multiple tissues (8). ATGL and its co-activator CGI-58 are central to a molecular complex including the perilipin family of proteins and G0S2 that orchestrate cLD catabolism in adipocytes as well as other tissue compartments.

Recent work in enterocytes indicates that the ATGL/CGI-58 pathway is active in regulating catabolism of enterocyte cLDs derived from the basolateral circulation but not those derived from the diet (9). Of note, these authors reported an increase in transcript of members of the Carboxylesterase (Ces) family of hydrolases in the intestine of *Atgl/Cgi-58* KO mice suggesting that these molecules may be involved in enterocyte cLD hydrolysis. The hydrolase(s) that regulate catabolism of cytoplasmic lipid droplets derived from dietary sources has not been identified.

We recently identified roles for the integrin ligand Milk Fat Globule Epidermal Growth Factor like 8 (MFGE8) and its receptor, the αvβ5 integrin, in intestinal lipid homeostasis. The MFGE8/integrin pathway links absorption of dietary fats with catabolism of small intestinal cLDs by promoting both enterocyte uptake of diet-derived luminal fats (10) and increased activity of enterocyte TG hydrolases resulting in TG mobilization from LDs for chylomicron production (11). Interestingly, unlike in the intestine, TG hydrolase activity is unaffected in white adipose tissue or liver of *Mfge8* KO mice (11) suggesting that enterocyte specific pathways regulate catabolism of diet-derived cLDs. In this work we investigated the molecular pathway through which MFGE8 regulates catabolism of enterocyte cLDs. We identify CES1D, a member of the Ces family of lipases, as the key hydrolase that functions downstream of MFGE8-integrin to mobilize fatty acids from cLD TG stores and regulate chylomicron production. We further show that dietary oleic acid increases expression and activity of CES enzymes through stabilizing protein levels of the transcription factor HNF4γ. The findings provide novel insight into intestinal regulation of postprandial lipid levels.

## RESULTS

### MFGE8 regulates the expression and activity of CES hydrolases

We have previously published that MFGE8 increases enterocyte TG hydrolase activity (11). To determine whether this effect is mediated through ATGL we isolated enterocytes from *Atgl* KO mice and assessed the effect of recombinant MFGE8 (rMfge8) on TG hydrolase activity. rMFGE8 significantly increased TG hydrolase activity in *Atgl* KO enterocytes and the effect size was similar to that of rMFGE8 on WT enterocyte TG hydrolase activity (supplementary figure 1) and *Mfge8* KO enterocyte TG activity (11). We interpret these data to indicate that the effect of MFGE8 on enterocyte TG hydrolase activity does not require ATGL.

We next took an unbiased approach to investigate which enterocyte TG hydrolases are regulated by MFGE8. We performed 3’ tag RNA sequencing of jejunal enterocytes isolated from WT and *Mfge8* KO mice and identified 530 differentially regulated genes (Figure 1A, Accession no. GSE200320). Ingenuity pathway analysis of these genes showed enrichment for triacylglycerol degradation related signaling (Figure 1B). Interestingly, we observed downregulation of several genes coding for hydrolases belonging to the CES1 family of enzymes in *Mfge8* KO mice (Figure 1C). We next performed activity-based staining (12) in WT and *Mfge8* KO intestinal cryosections using a fluorescently-labeled fluorophosphonate probe (TAMRA-FP) that binds the active confirmation of serine hydrolases. Cryosections from *Mfge8* KO mice showed markedly reduced fluorescence as compared with WT controls (Figure 1D) consistent with lower TG hydrolase activity. To further investigate which CES hydrolases had decreased activity in *Mfge8* KO enterocytes, we performed activity-based protein profiling (ABPP) with a serine hydrolase-specific fluorophosphonate biotin probe (FP-biotin) (13, 14). Consistent with our sequencing data, we found decreased activity for a subset of CES1 enzymes in *Mfge8* KO samples (Figure 1E, Accession no. MSV000089304). We interpret these data to indicate that MFGE8 regulates intestinal TG hydrolase activity through expression of CES family of enzymes.

**Figure 1:**
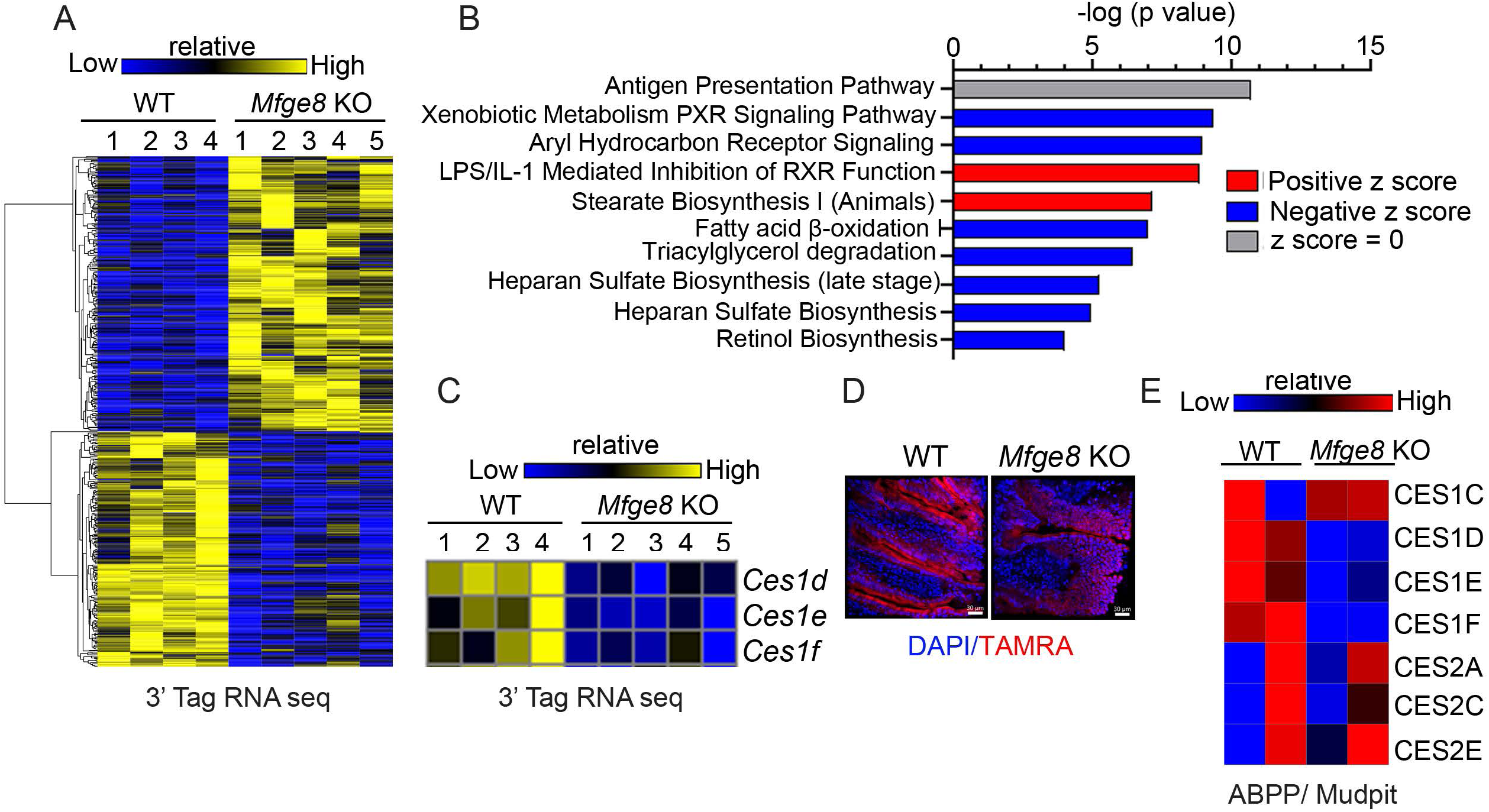
MFGE8 regulates the expression and activity of CES proteins. (A-C) 3’ Tag RNA sequencing of WT and *Mfge8* KO mouse primary enterocytes. (A) Heat map of differentially expressed genes. (B) Ingenuity pathway analysis of differentially expressed genes showing enriched biological processes. (C) Heat map showing altered expression of the Ces1 family genes. N=4 WT and 5 *Mfge8* KO 7-8 week-old male mice. (D) Confocal imaging of active serine hydrolases in small intestinal cryosections identified with a TAMRA-FP probe (red fluorescence) and counterstained with DAPI (blue). Representative image from 2 independent experiments, white bar = 30µM. (E) Serine hydrolase ABPP analysis showing differential activities of CES enzymes in WT and *Mfge8* KO primary enterocytes obtained from 8-9-week-old male mice. Each sample represents pooled enterocytes from 5 mice with LC/MS performed in technical duplicates.

### MFGE8 regulates the expression of CES hydrolases through the transcription factor HNF4γ

We next utilized the iRegulon database to identify putative candidate transcription factors that could mediate the effect of MFGE8 on *Ces* gene expression and cross-referenced these with transcription factor expression in WT enterocytes from our 3’ Tag RNA seq data. From this analysis, we found highest expression of the HNF4 family of transcription factors (that consist of Hnf4α and Hnf4γ) in WT enterocytes (Figure 2A). We subsequently analyzed available RNA sequencing data from a recent publication comparing gene expression of WT, *Hnf4γ* KO, and *Hnf4α* KO murine enterocytes (15). We found altered expression of multiple *Ces1* family genes in *Hnf4γ* KO (Figure 2B) but not in *Hnf4α* KO enterocytes (data not shown).

**Figure 2:**
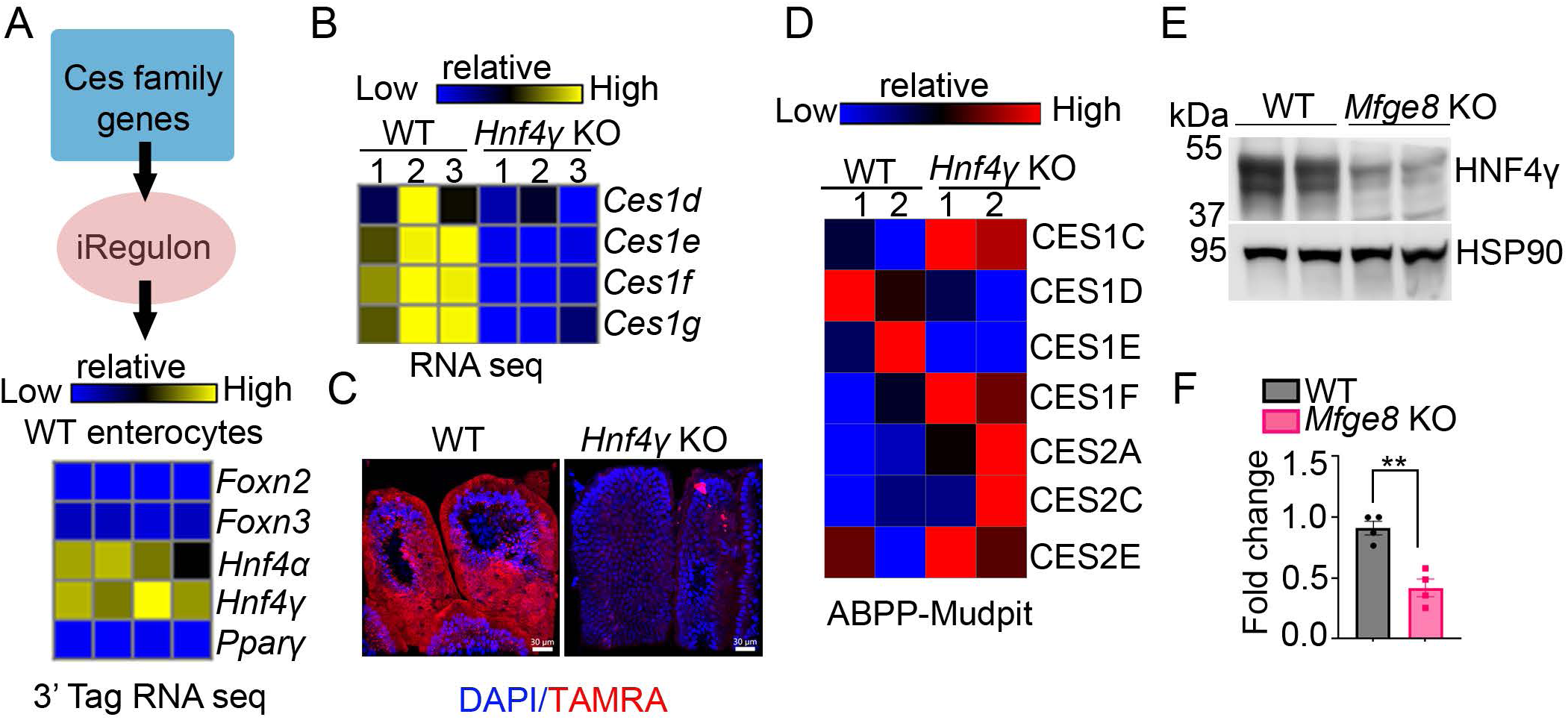
MFGE8 regulates the expression and activity of CES hydrolases through HNF4γ. (A) Heat map of expression of candidate transcription factors identified through the iRegulon database in 3’Tag RNA seq data of WT enterocytes. (B) Analysis of previously published RNA sequencing data (Accession no. GSE 200320) from WT and *Hnf4γ* KO enterocytes showing differential expression of the Ces genes. (C) Confocal imaging of active serine hydrolases identified with TAMRA-FP probe (red fluorescence) and counterstained with DAPI (blue) in WT and *Mfge8* KO intestinal cross-sections. Nuclei were stained with DAPI (blue). Representative image from 2 independent experiments, white bar = 30µM. (D) Serine hydrolase ABPP analysis showing differential activities of CES enzymes between WT and *Hnf4γ* KO primary enterocytes. N=2 independent experiments with each sample representing pooled enterocytes from 5 mice (total 10 mice per group). A mix of 9-10 week old male and female mice were used for this experiment. (E-F) Representative western blot of HNF4γ protein levels in WT and *Mfge8* KO enterocytes from 8-10-week-old male and female mice. Experiments are performed 2 independent times with a total of 4 mice in each genotype. (F) Densitometric analysis of the western blots (including panel E). All data expressed as Mean ± S.E.M. \*\**P* < 0.01. Data in panel F were analyzed by unpaired student’s t-test.

Next, we performed activity-based staining of serine hydrolases in WT and *Hnf4γ* KO intestinal cryosections and found a marked reduction in the hydrolase signal in the *Hnf4γ* KO group (Figure 2C). We also performed ABPP with FP-biotin and found that loss of HNF4γ led to reduced enzymatic activity of multiple CES1 subfamilies (Figure 2D, Accession no. MSV000089304) including CES1D. We next studied whether MFGE8 regulates HNF4γ transcript or protein expression. HNF4γ transcript was unchanged in *Mfge8* KO and WT jejunal enterocytes in our 3’ Tag RNA sequencing data set (Accession no. GSE200320). However, there was a marked reduction in HNF4γ protein levels in *Mfge8* KO enterocytes (Figure 2E-F). We interpret these data to indicate that MFGE8 modulates CES enzyme gene transcription by regulating HNF4γ protein levels.

### HNF4γ regulates catabolism of enterocyte cLDs

To investigate the functional role of HNF4γ in enterocyte LD homeostasis, we challenged WT and *Hnf4γ* KO mice with olive oil gavage (Figure 3A) and evaluated jejunal enterocyte TG hydrolase activity, small intestinal TG content, and serum TG levels. *Hnf4γ* KO enterocytes had significantly reduced TG hydrolase activity at baseline and 2 hours after olive oil gavage (Figure 3B). At 2 hours after the gavage, the increase in hydrolase activity associated with greater small intestinal TG content versus lower serum TG levels (figure 3C-E). We next administered 3H-labeled oleic acid by gavage to WT and *Hnf4γ* KO mice in the presence of the lipoprotein inhibitor tyloxapol (to prevent catabolism of serum TG) and measured the radioactive signal in the intestine and in the serum 2 hours later (Figure 3F-H). *Hnf4γ* KO mice had greater small intestinal radioisotope accumulation and reduced serum radiolabel (Figure 3G-H). The *Hnf4γ* KO mice were then fed a high-fat diet (HFD) or a control diet for 3 weeks. After a 12 hour fast, the *Hnf4γ* KO mice fed HFD had greater intestinal and lower serum TG content as compared with WT mice on a normal chow diet (Figure 3I,J). Of note, *Hnf4γ* KO mice exposed to acute or chronic fat challenges pheno-copied our previous findings with *Mfge8* KO mice (11) supporting the role of HNF4γ in catabolism of intestinal LDs (11).

**Figure 3:**
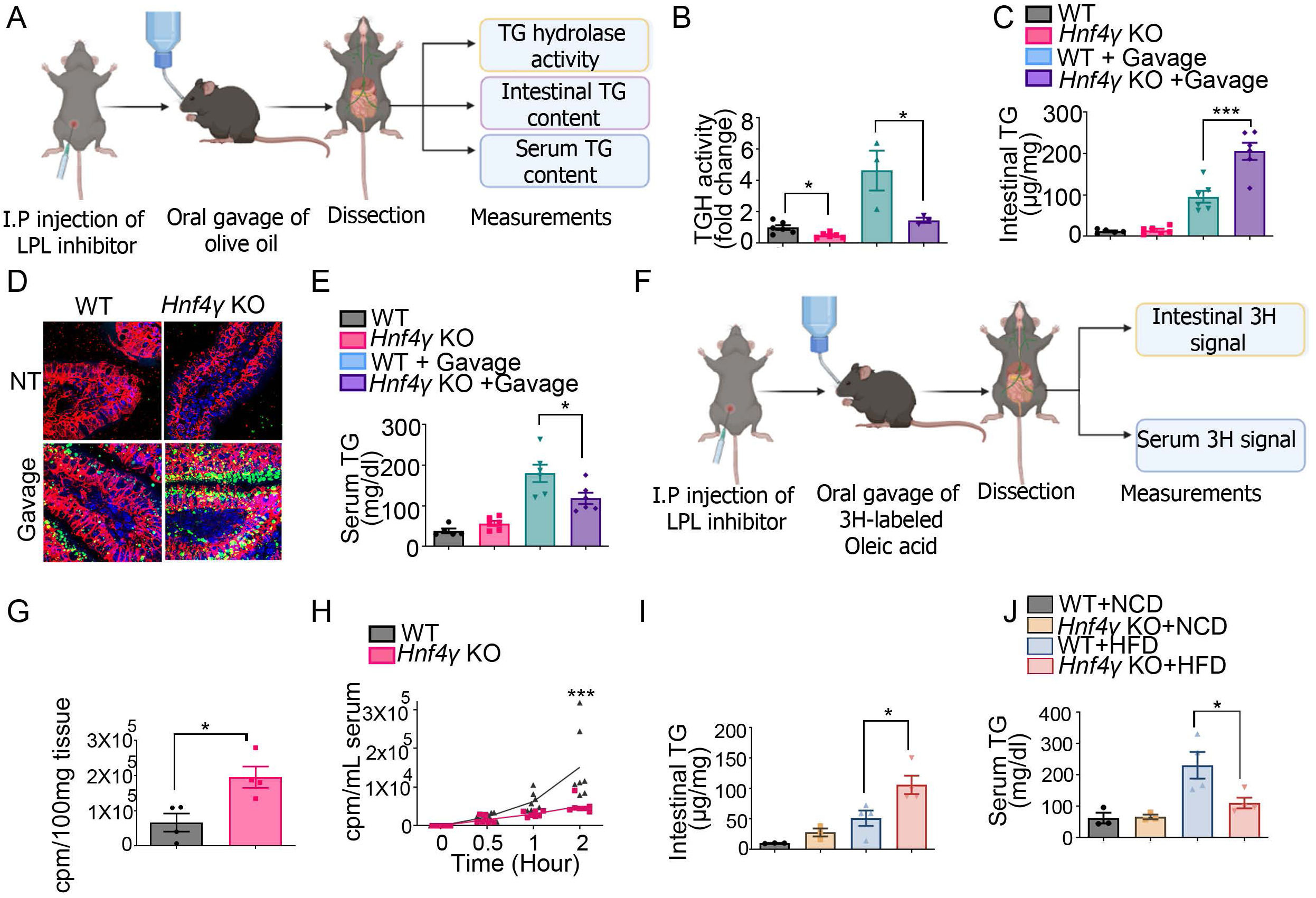
HNF4γ regulates catabolism of enterocyte cLDs. (A) Schema of the experimental design for panels B-E. (B) TG hydrolase (TGH) activity, (C, D) proximal jejunal TG content, and (E) serum TG content at baseline and 2 hours after olive oil gavage in primary enterocytes from WT and *Hnf4γ* KO mice (N=3-6 for panels B, C, E). (D) Intestinal tissue sections were stained with Bodipy (green) and anti-Epcam antibody (Magenta, N=2 in each group). Results are from 2 independent experiments. (F) Schema of the experimental design for panels G and H. (G) ^3^H signal in the proximal jejunum 2 hours after oral administration of ^3^H Oleic acid in WT and *Hnf4γ* KO mice. (H) ^3^H signal in the serum over time after oral administration of ^3^H Oleic acid. N=4 mice per group from 2 independent experiments. A mix of 7-10-week-old male and female mice were used for these experiments. (I-J) Intestinal (I) and serum (J) TG content of 8-week-old WT and *Hnf4γ* KO mice after 3 weeks on a HFD or normal-chow diet and following a 12 hour fast. N=3-4, data expressed as mean ± S.E.M. \**P* < 0.05, \*\**P* < 0.01, and \*\*\**P* < 0.001. Data in panels B, C, E, I and J analyzed by one-way ANOVA followed by Bonferroni’s posttest. Data in panel G analyzed by student’s t-test. Data in panel H were analyzed by 2-way ANOVA followed by Bonferroni’s posttest.

### CES1D regulates hydrolysis of enterocyte cLDs

We were next interested in understanding whether reduced expression of CES enzymes lead to impaired TG hydrolase activity in *Mfge8* KO and *Hnf4γ* KO enterocytes. The human genome contains 6 CES genes (*CES1, CES2, CES3, CES4A, CES5A*, and *CES1P1*). The mouse genome contains a larger number of CES proteins (20 have been annotated) due to tandem gene duplication (16). Of these, CES1d, CES1f, CES1g, CES2a, CES2b, CES2c and CES2e have known TG hydrolase activity (16-18). Expression of CES2 in the human intestine is well documented (19, 20). To delineate whether CES1 protein is expressed in human intestine, we performed Western blot of human small bowel epithelial cell lysates prepared from intestinal resections of patients with inflammatory bowel disease. Both CES1 and CES2 were expressed in these lysates (Figure 4A). Caco-2 cell lysates, a human colon carcinoma cell line known to express high levels of CES1 and low level of CES2 protein (21), were used as a positive control for these Western blots.

**Figure 4:**
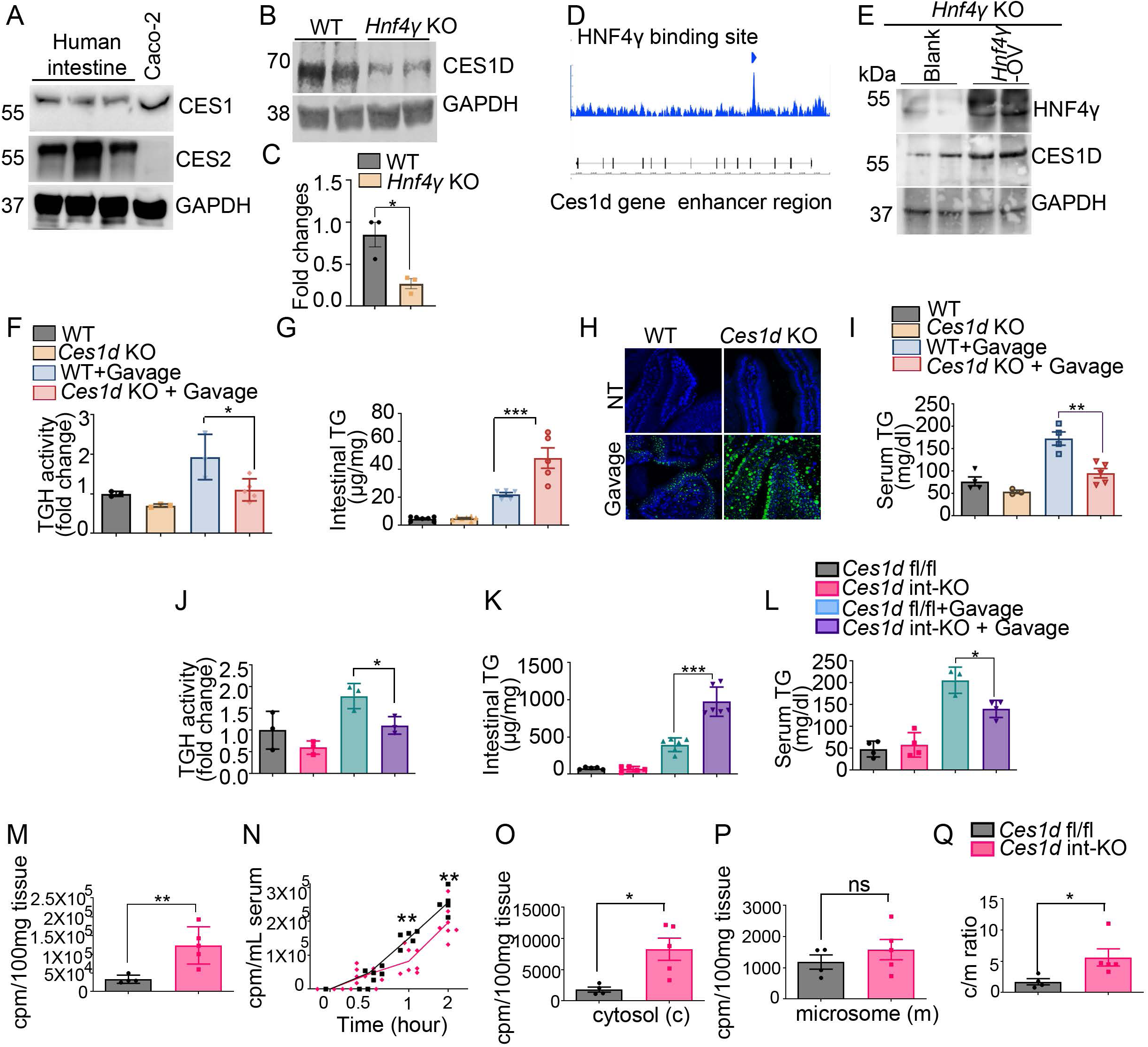
CES1D regulates hydrolysis of enterocyte cLDs. (A) Western blot of CES1 and CES2 in small intestinal lysates from human patients with inflammatory bowel disease. Caco-2 lysates serve as a positive control for CES1 expression with GAPDH a loading control. N=3 independent patient samples. (B) Representative western blot of CES1D in WT and *Hnf4γ* KO enterocyte lysates from 2 independent experiments, N=3 mice total. (C) Densitometric analysis of the western blots of CES1D (including panel B). (D) Analysis of previously published ChIP sequencing data (Accession no. GSE 112946) showing binding sites for HNF4γ on the promoter/enhancer regions of Ces family genes. (E) Western blot showing HNF4γ and CES1D protein level in *Hnf4γ* KO mice intestine segments incubated with Hnf4γ expressing or control adenovirus (Hnf4γ-AV). Western blot is representative of 3 independent experiments with N=4 *Hnf4γ* KO mice in total per experimental group. (F) Enterocyte TG hydrolase activity, (G,H) proximal jejunal TG content, and (E) serum TG content at baseline and 2 hours after olive oil gavage in WT and *Ces1d* KO mice. (H) Intestinal tissue sections from the same group of mice were stained with Bodipy (green). N=3-5 mice in each group. Results are from 2 independent experiments. (J) TG hydrolase activity in primary enterocytes, (K) TG content (K) in the proximal jejunum and (L) TG content in the serum at baseline and 2 hours after olive oil gavage in WT and *Ces1d* i-KO mice. N=3-6 mice in each group. Data merged from 2 independent experiments. (M) ^3^H signal in the proximal jejunjum 2 hours after oral administration of ^3^H Oleic acid. (N) ^3^H signal in the serum after oral administration of ^3^H Oleic acid over time in control and *Ces1d* int-KO mice. N=4-7 mice in each group. Results are from 2 independent experiments. (O-Q) ^3^H signal in the (O) cytosolic fraction, (P) microsomal fraction, and (Q) the ratio of cytosolic to microsomal radioactive signal in enterocytes from control and *Ces1d* int-KO mice 2 hours after oral gavage of ^3^H-labeled oleic acid. N=4-5 mice in each group. Data in panel F-G and panels I-L were analyzed by one-way ANOVA followed by Bonferroni’s posttest. Data in panels C, M, and O-Q were analyzed by student’s t-test. Data in panel N were analyzed by 2-way ANOVA followed by Bonferroni’s posttest. All data expressed as Mean ± S.E.M. \**P* < 0.05, \*\**P* < 0.01, and \*\*\**P* < 0.001.

We focused on CES1D since its expression (Figure 1C, 2B) and activity (Figure 1E, 2D) were significantly decreased in *Mfge8* and *Hnf4γ* KO enterocytes and because it is the murine ortholog of human CES1 (16). CES1D protein levels by Western blot were markedly reduced in *Hnf4γ* KO enterocytes (Figure 4B-C). We next analyzed data from recently published work looking at transcriptional targets of HNF4γ utilizing ChIP-sequencing in mouse enterocytes (16) and identified transcriptional binding sites for HNF4γ in the enhancer regions of *Ces1d* (Figure 4D). We next used an adenoviral vector to express exogenous Hnf4γ for 24 hours in in Hnf4γ KO intestines ex-vivo and subsequently probed by Western blot for HNF4γ and CES1D protein expression. Forced expression of HNF4γ in *Hnf4γ* KO enterocytes rescued CES1D protein levels (Figure 4E).

We then evaluated enterocyte LD homeostasis in *Ces1d* KO mice. Global *Ces1d* KO mice had reduced enterocyte TG hydrolase activity at baseline and after olive oil gavage (Figure 4F), which coupled with increased intestinal TG content (Figure 4G, H) and reduced serum TG levels (Figure 4I). Enterocyte-specific deletion of *Ces1d* (*Ces1d* int-KO using villin-Cre transgene) had significantly reduced enterocyte TG hydrolase activity at baseline and after olive oil gavage (Figure 4J) which associated with increased enterocyte TG content and reduced serum TG levels 2 hours after olive oil gavage (Figure 4K, L). Oral gavage of 3H-labeled oleic acid to *Ces1d* int-KO mice increased intestinal radioactivity (Figure 4M) and reduced it in serum as compared to controls (Figure 4N). Together, these data indicate that mice with global or intestine specific *Ces1d* deletion phenocopy *Mfge8* KO (11) and *Hnf4γ* KO mice in their response to olive oil gavage (Figure 3) in their impact on intestinal and serum lipids.

We have previously shown that *Mfge8* KO mice accumulate lipids in the cytosolic fraction after olive oil gavage consistent with altered cLD homeostasis (11). We therefore evaluated the intracellular location of accumulated TG in *Ces1d* int-KO mice 2 hours after 3H-labeled oleic acid gavage by fractionating jejunal enterocytes into cytosolic and microsomal components and measuring the radiolabel signal in each fraction. As with *Mfge8* KO mice (11), *Ces1d* int-KO mice accumulated radiolabel in the cytosolic fraction as compared with WT controls with no apparent differences in the microsomal fraction (Figure 4O-Q). We confirmed the relative enrichment of cytosolic and microsomal fractions by western blotting for cytosolic marker protein GAPDH and microsomal marker protein BIP (Supplementary Figure 2).

### MFGE8 regulates TG hydrolase activity through CES1D

To determine if MFGE8 and the αvβ5 integrin modulate CES1D protein levels we performed Western blot in *Mfge8* and *β5* KO enterocytes and found marked reduction of CES1D in both populations (Figure 5A-D). To directly assess whether the effect of MFGE8 on cLD catabolism is mediated through CES1D, we evaluated the ability of rMFGE8 to increase TG hydrolase activity in *Ces1d* KO enterocytes. While rMFGE8 significantly increased TG hydrolase activity in WT and *Mfge8* KO enterocytes, it had no effect in *Ces1d* KO enterocytes (Figure 5E). We used a mutated Mfge8 protein construct (RGE) that cannot bind integrins (10) as a negative control (Figure 5E). We next assessed whether transgenic re-expression of MFGE8 specifically in enterocytes in *Mfge8* KO mice using a tetracycline-inducible system (22) modulated CES1D protein levels. Inducible expression of MFGE8 rescued the loss of CES1D protein levels (Figure 5F-G) as well as the TG hydrolase activity (Figure 5H). We then assessed cLD catabolism in mice with global deletion of both *Ces1d* and *Mfge8* (*Ces1d/Mfge8* KO). We administered 3H-labeled oleic acid by gavage to WT, *Ces1d* KO, *Mfge8* KO, and *Ces1d/Mfge8* KO mice and quantified 3H radiolabel in the small intestine and serum 2 hours after gavage (Figure 5I-J). *Ces1d/Mfge8* KO mice had a similar increase in intestinal radiolabel and a similar reduction in serum radiolabel as *Mfge8* and *Ces1d* KO mice indicating the loss of both alleles had no additive effect (Figure 5I-J). Together, these data indicate that MFGE8 modulates enterocyte TG hydrolase activity in large part through CES1D.

**Figure 5:**
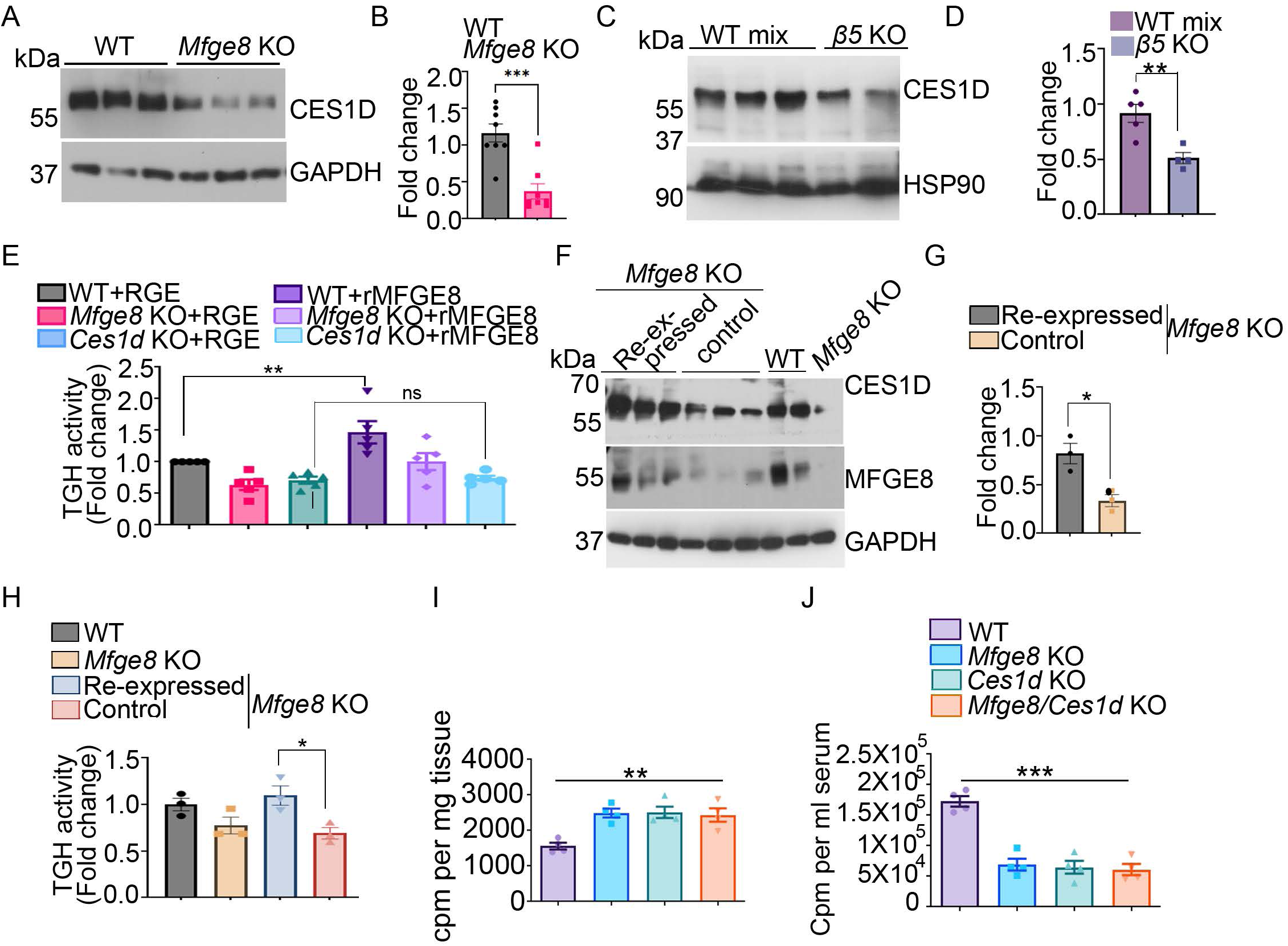
MFGE8 regulates TG hydrolase activity through CES1D. (A-B) Representative western blot (A) showing CES1D protein level in WT and *Mfge8* KO primary enterocytes from 3 independent experiments. GAPDH was used as loading control. N=9 mice per group. (B) Densitometric analysis of the western blots (including panel A). (C) Representative western blot showing CES1D protein level in WT and *β5* KO primary enterocytes from 2 independent experiments. HSP90 was used as loading control. N=5 WT and 4 *β5* KO mice in total. (D) Densitometric analysis of the western blots of CES1D protein (including panel C). (E) TG hydrolase activity in WT, *Ces1d* KO and *Mfge8* KO primary enterocytes 1 hour after incubation with rMFGE8 or RGE. N=5 independent experiments. (F) Western blot of CES1D and MFGE8 protein levels in enterocytes of *Mfge8* KO mice with transgenic inducible expression of MFGE8 in enterocytes (MFGE8 reexpressed, Vil rtTA+ TetO Mfge8+) and single transgenic, WT, and *Mfge8* KO enterocyte controls. GAPDH was used as loading control. (G) Densitometric analysis of the blot presented in panels F. (H) TG hydrolase activity in primary enterocytes isolated from the same groups of mice in panel E-F. N=3 mice in each group. (I-J) ^3^H signal in the proximal jejunjum (I) and serum (J) 2 hours after oral administration of ^3^H Oleic acid to WT, *Mfge8* KO, *Ces1d* KO and *Mfge8/Ces1d* double KO mice. N=3-4 mice in each group. All data expressed as Mean ± S.E.M. \**P* < 0.05, \*\**P* < 0.01, and \*\*\**P* < 0.001. Data in panels B, D and G were analyzed by unpaired t-test. Data in panels E and H through J were analyzed by one-way ANOVA followed by Bonferroni’s posttest.

### MFGE8 links fatty acid absorption to LD catabolism through HNF4γ

We have previously shown that MFGE8 promotes absorption of dietary fatty acids in the small intestine. HNF4γ is a nuclear hormone receptor that constitutively binds saturated and cis-monounsaturated fatty acids of 14-18 carbons (23). We therefore examined whether MFGE8-mediated fatty acid absorption impacts the activity of HNF4γ. In our 3’ Tag RNA seq data (Accession no. GSE200320) we found in *Mfge8* KO enterocytes decreased expression of *Cd36* and *Fatp2* (Figure 6A), two fatty acid transporters active in the intestine (24-27). We therefore assessed whether genetic deletion of *Cd36* impacts HNF4γ and observed a marked reduction in HNF4γ protein level in *Cd36* KO enterocytes (Figure 6B,C). Moreover, genetic deletion of *Cd36* also caused a reduction in CES1D protein level in enterocytes (Figure 6B,C).

**Figure 6:**
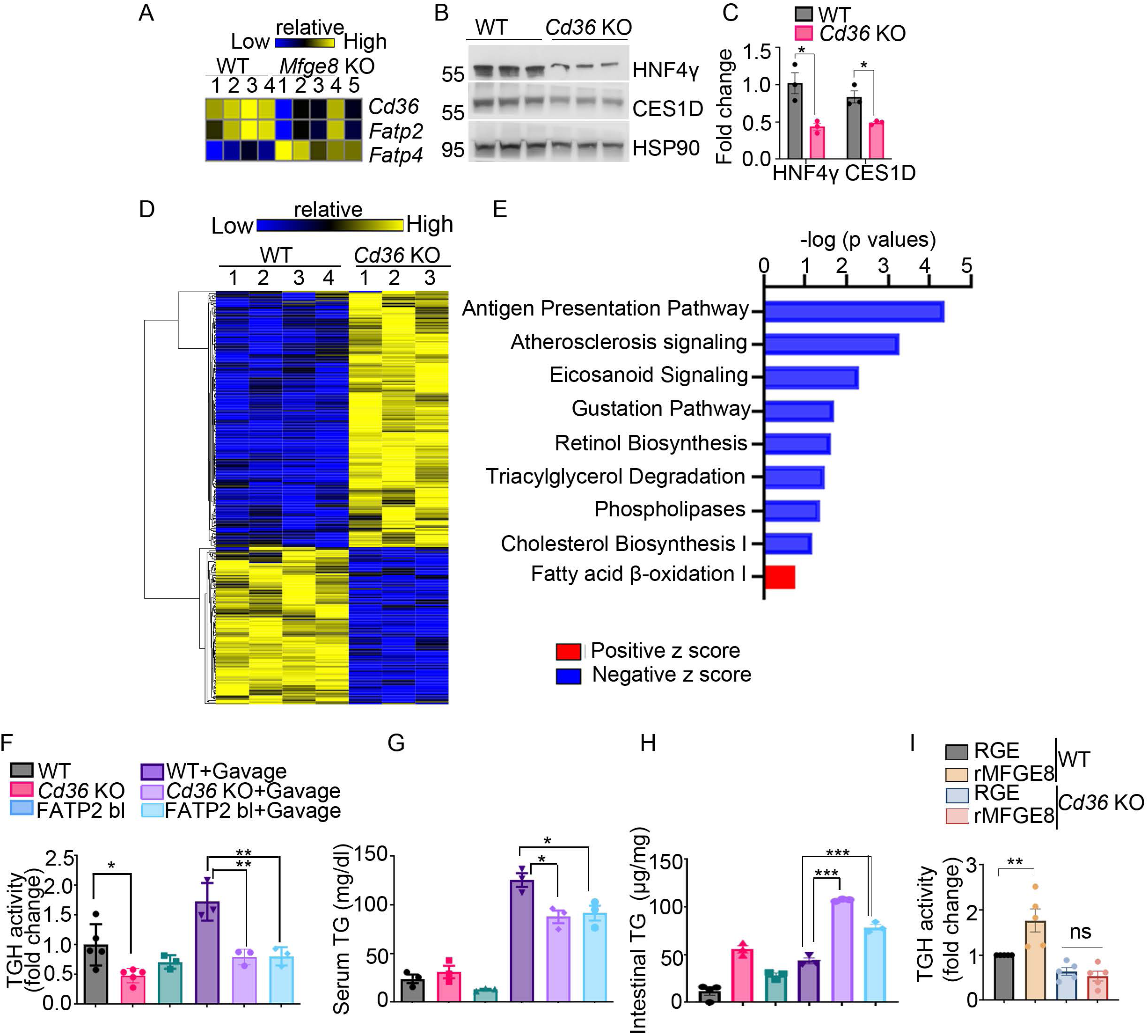
MFGE8 links fatty acid absorption to LD catabolism through HNF4γ. (A) Heatmap showing differentially expressed fatty acid transporters in WT and *Mfge8* KO primary enterocytes from 3’ Tag RNA sequencing (Accession: GSE 200320). (B-C) Western blot of HNF4γ and CES1D protein level in WT and *Cd36* KO primary enterocytes. N=3 mice in each group. (C) Densitometric analysis of the western blot in panel B. (D-E) Heatmap (D) and ingenuity pathway analysis (E) of differentially expressed genes in 3’ Tag RNA sequencing of WT and *Cd36* KO intestine. (F-H) Primary enterocyte (F) TG hydrolase activity, (G) TG content in the proximal jejunum, and (H) serum TG content at baseline and 2 hours after acute fat challenge of WT, *Cd36* KO and WT mice treated with pharmacological inhibitor of FATP2 (FATP2 bl). (I) TG hydrolase activity in WT and *Cd36* KO primary enterocytes 1 hour after incubation with rMFGE8 or RGE. N=5 independent experiments. Data expressed as Mean ± S.E.M. \**P* < 0.05, \*\**P* < 0.01, and \*\*\**P* < 0.001. Data in panels C were analyzed by unpaired t-test. Data in panels F through I were analyzed by one-way ANOVA followed by Bonferroni’s posttest.

We next performed 3’Tag-RNA seq of WT and *Cd36* KO small intestine (Figure 6D). Ingenuity pathway analyses of differentially expressed genes indicated enrichment of triglyceride degradation processes (Figure 6E). Furthermore, TG hydrolase activity was significantly decreased in *Cd36* KO enterocytes at baseline and after acute fat challenge (Figure 6F). Pharmacological blockade of FATP2 in WT mice also suppressed enterocyte TG hydrolase activity after acute fat challenge (Figure 6F). Both *Cd36* KO mice and WT mice treated with a pharmacological inhibitor of FATP2 accumulated lipids in the small intestine and had lower TG level after an acute fat challenge as compared with WT controls (Figure 6 G,H). To further assess whether the effect of MFGE8 on LD catabolism involves CD36, we evaluated the ability of rMFGE8 to increase TG hydrolase activity in *Cd36 KO* enterocytes. While rMFGE8 significantly increased TG hydrolase activity in WT enterocytes, it had no effect in Cd36 KO enterocytes (figure 6I). These data suggest that the effects of MFGE8 on enterocyte HNF4γ protein levels and LD catabolism are linked through MFGE8/CD36 dependent fatty acid absorption.

### Fatty acid stabilizes HNF4γ protein to activate transcription of Ces genes

We next assessed whether oleic acid activates HNF4γ-mediated transcription of *Ces* genes associated with lipid catabolism. We cloned the 500-bp region of the putative enhancer regions of *Ces1d* into a luciferase vector and performed a dual luciferase activity assay in control and HEK293 cells overexpressing HNF4γ (via adenovirus) in presence and absence of oleic acid (Figure 7A). Cells with HNF4γ overexpression had significantly increased luciferase activity for *Ces1d*. Interestingly, oleic acid further induced transcription of *Ces1d* (Figure 7B). We next performed a 24-hour cycloheximide pulse-chase experiment in HEK293 cells in which we overexpressed HNF4γ by adenovirus, and subsequently incubated cells with oleic acid (Figure 7C). HNF4γ protein levels decreased at the 12-hour time point in presence of cycloheximide but addition of oleic acid prevented this decay (Figure 7D-E). We interpret these data to indicate that oleic acid induces *Ces1d* transcription by stabilizing enterocyte HNF4γ protein levels. We then determined whether a chronic HFD impacts expression of HNF4γ and CES1D in vivo. After 3 weeks on HFD, small intestinal protein levels of HNF4γ and CES1D were increased in mice on HFD as compared with normal chow diet (Figure 7F-G). We interpret these data to indicate that stabilization of HNF4γ protein levels by dietary fatty acids drives the increase in CES1D expression.

**Figure 7:**
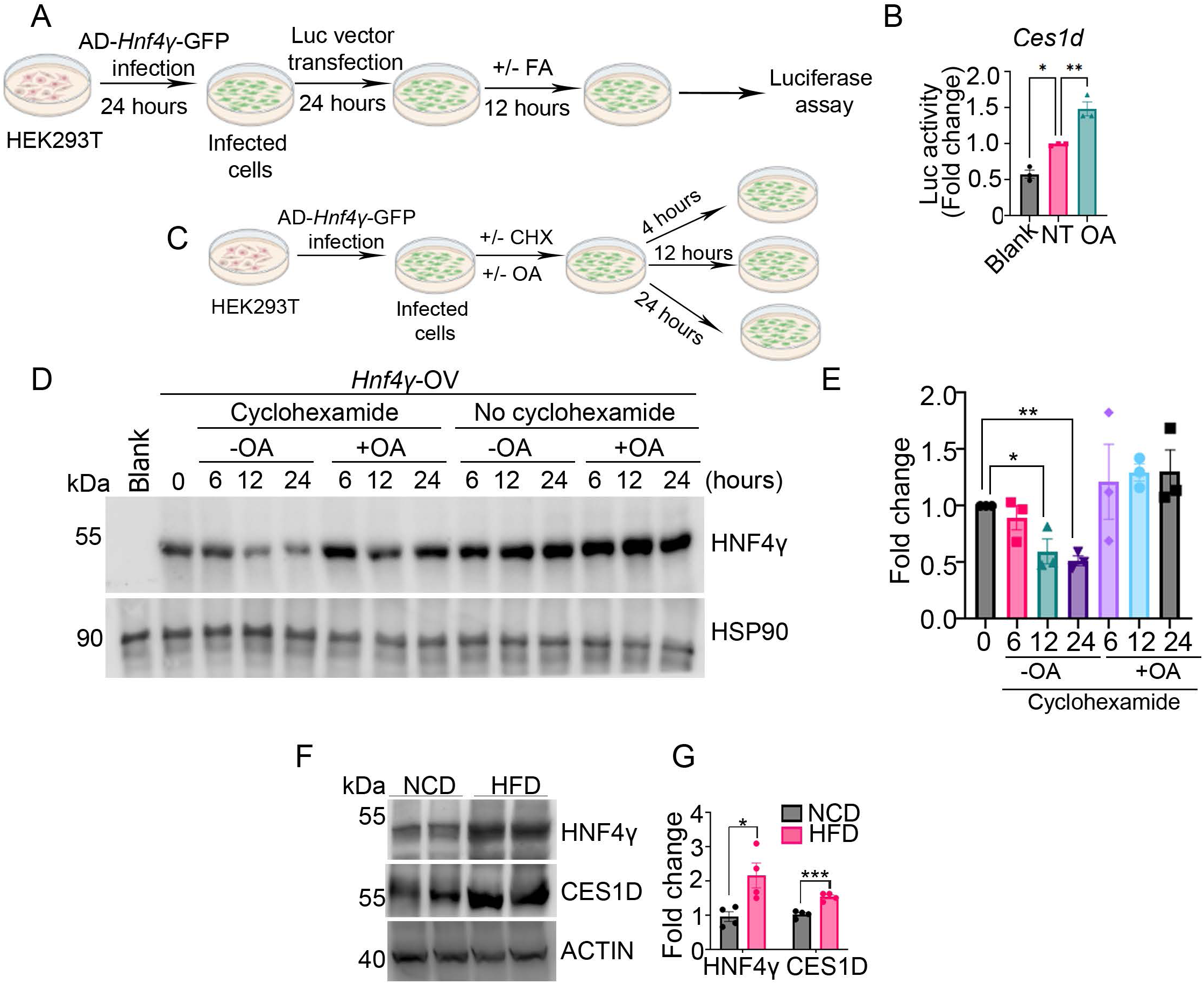
Fatty acid stabilizes HNF4γ protein to activate transcription of Ces genes. (A) Schematic representation of the method used for dual luciferase assay presented in panel B. (B) Data showing normalized luciferase activity of the Ces1d gene enhancer in the presence of oleic acid in HEK293 cells with adenoviral overexpression of Hnf4γ. Cells infected with a blank adenovirus (blank) used as a negative control. N=3 independent experiments. (C) Schematic representation of the method used for the cyclohexamide chase assay presented in panel D. (D) Representative western blot showing HNF4γ protein levels 6, 12 and 24 hours after treatment with oleic acid or DMSO control in the presence and absence of cyclohexamide in HEK 293 cells. Western blot is representative of 3 independent experiments. (E) Densitometric analysis of HNF4γ protein levels (including panel D). (F) Representative western blot showing HNF4γ and CES1D protein level in the small intestine of mice on a normal chow or high-fat chow diet. Results are from 2 independent experiments. (G) Densitometric analysis of the western blots of HNF4γ and CES1D (including panel F). N=2 mice per group per experiment (Total 4 mice in each group). All data expressed as Mean ± S.E.M. \**P* < 0.05, \*\**P* < 0.01. Data in panels B and E were analyzed by one-way ANOVA followed by Bonferroni’s posttest. Data in panel G were analyzed using an unpaired student’s t-test.

### The effect of MFGE8 of cLD catabolism is unique to diet-derived cLDs

To determine whether MFGE8 modulates cLDs derived from the basolateral circulation and not from luminal fat absorption we administered ^3^H-oleic acid IP to *Mfge8* KO and WT mice and quantified the radioactive signal in the small intestine. Interestingly, we did not observe differences when comparing *Mfge8* KO and WT samples (Supplementary Figure 3A). Furthermore, when we performed cell fractionation and measured the radioactive signal in cytosolic and microsomal fractions, we found no significant differences between *Mfge8* KO and WT samples. (Supplementary Figure 3B-D). These data support the interpretation that the effect of MFGE8 on cLD hydrolysis is restricted to diet-derived cLDs.

### β5 blockade impairs hydrolysis of enterocyte cLDs

Finally, we determined the therapeutic potential of β5 integrin blockade on dampening postprandial lipemia. IP β5 blockade reduced TG hydrolase activity and increased small intestine lipid accumulation after an acute olive oil gavage as compared with isotype control antibody in WT mice (Figure 8A-B).

**Figure 8:**
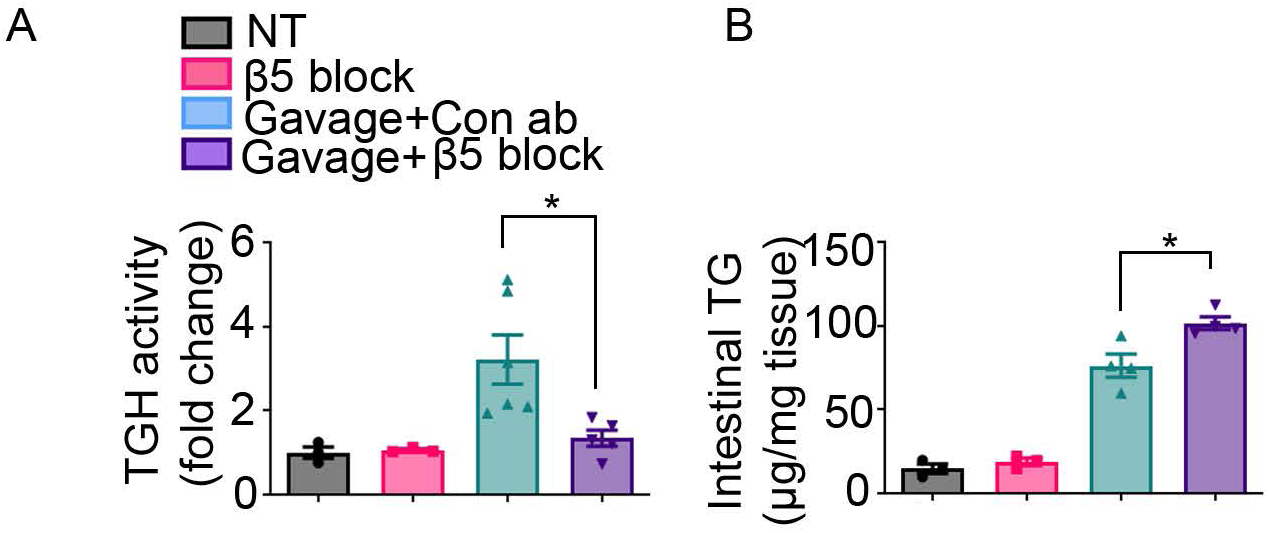
β5 blockade impairs hydrolysis of enterocyte cLDs. (A,B) TG hydrolase activity (A) in primary enterocytes and proximal jejunal TG content (B) at baseline and 2 hours after acute fat challenge in WT mice treated with either β5 blocking antibody or control antibody. N=3-5 mice in each group. Data expressed as Mean ± S.E.M. \**P* < 0.05. Data were analyzed by one-way ANOVA followed by Bonferroni’s posttest.

**Figure 9:** Schematic representation showing how MFGE8-mediated fatty acid uptake activates HNF4γ-mediated transcription of Ces genes leading to postprandial lipemia.

## Discussion

Enterocytes are unique in that they are polarized cells that absorb fatty acids from 2 distinct cellular pools: circulating fatty acids from the basolateral surface and dietary fatty acids from the apical surface. Absorbed fatty acids can be catabolized through beta-oxidation, packaged into chylomicrons for delivery through the circulation to peripheral organs, or retained in the enterocyte as part of cLDs. Storage of fatty acids in cLDs modulates the risk of developing atherosclerotic disease by minimizing the extent of postprandial lipemia, particularly in the setting of a fat-rich diet. The importance of this regulatory mechanism is evident when one considers that humans with obesity and/or diabetes characteristically have exaggerated postprandial lipemia (28, 29) coupled with a marked increase in the risk of developing coronary artery disease. Of note, oxidation of chylomicron remnants is particularly pro-atherogenic (30, 31) providing a rationale for why serum lipid levels after a meal have a stronger correlation with coronary artery disease than fasting serum lipid levels (2).

Enterocytes have 2 distinct pools of cLDs; those derived from lipids from the circulation and those from lipids absorbed from the diet (5, 32). CLDs derived from the circulation are thought to be primarily utilized for phospholipid synthesis or beta-oxidation while those from the diet are thought to be primarily incorporated into TGs used for chylomicron production (5, 32). Catabolism of each cLD pool occurs through distinct molecular pathways with hydrolysis of enterocyte cLDs derived from the circulation occurring by the same molecular pathways utilized by adipocytes and centered on the ATGL/CGI-58 (7-9, 33).

We have previously shown that MFGE8 links the absorption of dietary fats with mobilization of fatty acids from cLDs for chylomicron production through ligation of αv integrins (10, 11). We were therefore interested in understanding which TG hydrolases function downstream of the MFGE8-integrin pathway. Our findings that recombinant MFGE8 significantly increased enterocyte TG hydrolase activity in *Atgl* KO enterocytes indicated that the effect of the MFGE8-integrin axis on cLDs occurs through an unidentified TG hydrolase. Using an unbiased approach, we found differential expression of the CES enzyme family of hydrolases in *Mfge8* KO enterocytes and subsequently show that one member, CES1D, mediates the bulk of effect of the MFGE8-integrin axis on enterocyte CLD homeostasis.

Of note, while the human genome contains 6 CES genes (*CES1, CES2, CES3, CES4A, CES5A*, and *CES1P1*), tandem gene duplication has led to 20 annotated Ces enzymes in the mouse genome (16). The CES1 family in mice consists of 8 members, 3 of which had decreased expression in *Mfge8* KO enterocytes (*Ces1d*-*f*). We chose to focus on CES1D because it is the closest murine ortholog of human *CES1*. Whether CES1D mediates the entirety of the effect of MFGE8 on enterocyte TG hydrolase activity is difficult to ascertain given the number of CES genes that have altered expression or activity in *Mfge8* KO enterocytes. However, it is clear from the work presented here that CES1D mediates the majority, if not all, of the effects of MFGE8 on enterocyte TG hydrolysis given how closely the *Ces1d* KO mice phenocopy *Mfge8* KO mice in their response to acute and chronic fat challenges, the failure of recombinant MFGE8 to increase TG hydrolase activity in *Ces1d* KO enterocytes, and the lack of additive effects on enterocyte TG content and serum TG levels after olive oil gavage in mice double KO for *Mfge8* and *Ces1d* (as compared with single KO mice).

One potential limitation is that genetic deletion of *Ces1d* altered expression of other *Ces* enzymes, which could have contributed to the observed physiological effects. Our study focused on CES1D, as the closest murine ortholog of the human hydrolase CES1 and as an enzyme shown to express and have high activity in the proximal intestine (34) where most chylomicron generation is known to occur. A profiling of the enzymatic activity of hydrolases in the murine small intestine identified multiple CES enzymes, including CES1D, and highlighted the need to understand the function and regulation of the various intestinal hydrolases. Our data suggest that CES1D might be a rate-limiting enzyme in catabolism of LDs from dietary lipids, such that its deletion could impact the overall pathway, despite potential involvement of other CES enzymes. Of note, increased transcription of *Ces* enzymes in the intestine of mice with enterocyte-specific deletion of ATGL and CGI-58 has been reported (9) suggesting that some Ces enzymes might compensate for loss of the canonical pathway for hydrolysis of LDs derived from the circulation.

The effect of the Mfge8-integrin pathway being restricted to diet-derived cLDs is consistent with our published work showing no differences in TG hydrolase activity in the liver or white adipose tissue of *Mfge8* KO mice (11). We conclude from the work presented here that the CES family of enzymes, in particular CES1D, mediate the effect of MFGE8 on diet-derived LDs based on several observations. First, recombinant MFGE8 retains the ability to increase enterocyte TG hydrolysis in enterocytes KO for *Atgl*, the enzyme that regulates ctabolism of LDs derived from the basolateral circulation (9). Second, gavage of radiolabeled oleic acid in the setting of pretreatment (for the *Ces1d* KO and *HNFγ* KO studies) with the lipoprotein lipase inhibitor Tyloxapol (which prevents breakdown of serum TG that is a prerequisite for absorption of fatty acids from the basolateral circulation) leads to accumulation of radiolabel in cLDs in the small intestine of *Mfge8* KO (11), *Ces1d* KO, and *HNFγ* KO mice. However, IP administration of 3H-oleic acid to *Mfge8* KO and WT mice leads to similar small intestine total, cytosomal, and microsomal fraction radioactive signals arguing against an effect of MFGE8 on absorption of fatty acids from the basolateral surface and accumulation of basolateral-derived LDs (that are present in the cytosolic fraction).

Our previous work identifies a biological program linking absorption of dietary fat with mobilization of fat stored in enterocyte LDs for chylomicron production through MFGE8 (9-11). Here, we further delineate the molecular mechanisms coupling these two processes by showing that a dietary fatty acid (oleic acid), absorbed in part through MFGE8-dependent mechanisms, stabilizes protein levels and transcriptional activity of a nuclear hormone receptor, HNFγ, which then increases enterocyte CES enzyme expression and cLD hydrolysis. Both members of the HNF family of transcription factors, HNFα and HNFγ, have been shown to constitutively bind fatty acids (23). HNFγ expression is predominately restricted to the small intestine (15). To explore the hypothesis that fatty acid uptake is an important step in MFGE8-induced increases in CES expression/activity, we focused on CD36, a fatty acid transporter with a well-established role in absorption of fatty acids in the proximal intestine (35). Furthermore, we had previously shown that in adipocytes and hepatocytes, MFGE8 induces cell surface translocation of CD36 leading to enhanced fatty acid uptake (10). Interestingly, here we found markedly decreased expression of Cd36 in *Mfge8* KO enterocytes. Furthermore, *Cd36* KO enterocytes phenocopied *Mfge8* KO enterocytes with respect to HNFγ and CES1D protein expression, TG hydrolase activity, and differential gene expression profiles in enterocytes. Additionally, recombinant MFGE8 failed to increase TG hydrolase activity in *Cd36* KO enterocytes.

These data suggesting that fatty acid uptake regulates HNFγ-dependent transcription were further supported by showing positive regulation by oleic acid of HNFγ-dependent CES transcription and HNFγ protein levels. In sum, these data indicate to us that MFGE8-CD36 dependent uptake of dietary fats promotes enterocyte TG hydrolase activity by stabilizing HNFγ-leading to increased HNFγ-dependent transcription of CES enzymes. Our findings generate new questions related to the specific diet-derived fatty acids and/or metabolites that serve as HNF4γ ligand, whether these fatty acids replace the constitutively bound fatty acid in HNFγ and how this interplay regulates the transcriptional activity of HNF4γ.

The MFGE8-integrin pathway has emerged as an interesting candidate for therapeutic targeting in metabolism. We have previously shown that MFGE8 promotes the development of obesity both through a direct effect on intestinal fat absorption (10) and by reducing gastrointestinal motility thereby allowing more time for nutrient absorption (22). Furthermore, these effects can be therapeutically targeted independent of each other given that they are mediated by different integrin receptors (αvβ5 for fat absorption and α8β1 for motility effects). More recently, we have shown that MFGE8 ligation of αvβ5 induces insulin resistance at the level of the insulin receptor and that blockade of this pathway leads to enhanced insulin sensitivity in the skeletal muscle and liver (36). Our work here identifies a carboxylesterase enzyme that is responsible for the effect of MFGE8 on catabolism of diet-derived cLDs. This pathway can be targeted to reduce the severity of postprandial lipemia in obese, insulin-resistant patients while concurrently reducing fat absorption (10) and enhancing peripheral tissue insulin sensitivity (36). Whether these benefits would outweigh the potential risks of targeting this biological pathway remains to be determined.

## Materials and methods

### Mice

All animal experiments were approved by the UCSF Institutional Animal Care and Use Committee in adherence to NIH guidelines and policies. *Mfge8* KO mice were purchased from RIKEN and are in the C57BL/6 background and have been extensively characterized. *Ces1d* KO and *Ces1d* flox/flox have been previously characterized (37-39). *Villin-Cre* transgenic mice [Tg (Vil1 Cre) 997Gum)] (Jackson laboratories) were bred with *Ces1d* flox/flox mice to generate intestine-specific Ces1d KO (*Ces1d* int-KO) mice (*Ces1d* flox/flox Vil Cre+). *Ces1d* flox/flox Vil Cre negative mice were used as control. *Hnf4α* flox/flox Vil Cre ert2 / *Hnf4γ*^Crispr^ mice are in a mixed background and have been characterized (15). We described Hnf4α flox/flox Vil Cre ert2 Hnf4γ^Crispr^ mice as *Hnf4γ* KO for our experiments. *Hnf4α* flox/flox Vil Cre ert2 *Hnf4γ*+/+ mice were used as controls. Tg(TetO-Mfge8) transgenic mice containing the tetracycline-inducible Mfge8 construct were crossed with a *Mfge8* KO mice line created using a gene disruption vector (40) and mice carrying the Tg(Vil-rtTA) transgene. *Cd36* KO mice has been extensively characterized (41).

### Isolation of primary enterocytes

Primary enterocytes were harvested from intestinal jejunal segments following previously published protocol (11). The jejunal lumen was washed with buffer A (115 mM NaCl, 5.4 mM KCl, 0.96 mM NaH2PO4, 26.19 mM NaHCO3, and 5.5 mM glucose buffer at pH 7.4, gassed for 30 minutes with 95% O2 and 5% CO2) and subsequently filled with buffer B (67.5 mM NaCl, 1.5 mM KCl, 0.96 mM NaH_2_PO_4_, 26.19 mM NaHCO3, 27 mM sodium citrate, and 5.5 mM glucose at pH 7.4, saturated with 95% O_2_ and 5% CO_2_) and incubated in buffer B for 15 minutes at 37 with constant shaking. After 15 minutes, the solution was discarded and the jejunal segments were transferred to a new 100 mm dish and filled with and incubated in buffer C (115 mM NaCl, 5.4 mM KCl, 0.96 mM NaH_2_PO_4_, 26.19 mM NaHCO_3_, 1.5 mM EDTA, 0.5 mM dithiothreitol, and 5.5 mM glucose at pH 7.4, saturated with 95% O_2_ and 5% CO_2_) for 15 minutes at 37 with constant shaking after which the luminal contents were centrifuged and the pellet containing the epithelial cells used for subsequent experiments. The purity of the isolation was checks by FACS sorting cells using anti-Epcam antibody. Epcam-positive cells constituted 85-90% of the isolated cell pellet.

### Recombinant MFGE8 (rMFGE8)

rMFGE8 and RGE constructs consisted of murine cDNA of *Mfge8* (long isoform) fused with the human FC domain. They were expressed in High Five cells and affinity purified as previously described (10, 36). The RGE construct contains a point mutation that changes the integrin-binding RGD sequence to RGE. Primary enterocytes were treated with either RGE or rMFGE8 (10 mg/ml) for 1 hour and then washed in PBS and processed for TG hydrolase activity assay.

### TG hydrolase (TGH) activity assay

TG hydrolase activity from jejunal enterocytes was measured using previously a published protocol (11). Protein was extracted from primary enterocytes in 100 mM potassium phosphate buffer by brief sonication. For Figure 5A, primary enterocytes were incubated with rMFGE8 or RGE proteins (10 µg/mL) in serum-free media for 1 hour before proceeding with protein isolation. 60-100 µg protein was incubated with 100 μl TG substrate (25 nmol triolein/assay and 40,000 cpm/nmol ^14^C-triolein; PerkinElmer) and 35.5 μg mixed micelles of phosphatidylcholine and phosphatidylinositol (3:1, w/w), respectively, for 1 hour at 37. After 1 hour, the reaction was terminated by adding 3.25 ml methanol/chloroform/heptane (10:9:7, v/v/v) and 1 ml 100 mM potassium carbonate (pH 10.5 with boric acid). After centrifugation (800×g, 15 minutes, 4°C), radioactivity was measured in 1 ml of the upper phase by liquid scintillation counting. The radioactivity counts were normalized relative to protein concentration and the TG hydrolase activity was expressed as relative fold changes to the untreated control samples.

### RNA isolation

RNA from primary enterocytes was isolated using Qiagen RNeasy plus micro kit. RNA from small intestinal tissues was isolated using Qiagen RNeasy lipid tissue mini kit.

### 3’ Tag RNA sequencing

Gene expression profiling of primary enterocyte RNA samples and total intestinal RNA samples were carried out using a 3’-Tag-RNA-Seq protocol. Barcoded sequencing libraries were prepared using the QuantSeq FWD kit (Lexogen, Vienna, Austria) for multiplexed sequencing according to the recommendations of the manufacturer using the UDI-adapter and UMI Second-Strand Synthesis modules (Lexogen). High integrity total RNA samples were processed according to the QuantSeq default protocol. The fragment size distribution of the libraries was verified via micro-capillary gel electrophoresis on a LabChip GX system (PerkinElmer, Waltham, MA). The libraries were quantified by fluorometry on a Qubit fluorometer (LifeTechnologies, Carlsbad, CA), and pooled in equimolar ratios. The library pool was Exonuclease VII (NEB, Ipswich, MA) treated, SPRI-bead purified with KapaPure beads (Kapa Biosystems / Roche, Basel, Switzerland), quantified via qPCR with a Kapa Library Quant kit (Kapa Biosystems) on a QuantStudio 5 RT-PCR system (Applied Biosystems, Foster City, CA). Up to 48 libraries were sequenced per lane on a HiSeq 4000 sequencer (Illumina, San Diego, CA) with single-end 100 bp reads.

### Bioinformatic analysis

FASTQ files were trimmed with Trimmomatic v0.38.1 (https://github.com/usadellab/Trimmomatic) and umi-tools (https://github.com/CGATOxford/UMI-tools) in order to remove low quality reads and any adapter contamination. The reads were mapped with HISAT2 v2.1.0 to the mouse genome (mm10 / GRCm38). After mapping, all BAM files were used as input for HTSeq-count v0.91 to calculate transcript coverage. DESeq2 (v2.11.40) (42) was used to find differentially expressed transcripts between samples for each sequencing depth. Differentially expressed genes (DEG) were determined based on whether the adjusted FDR was ≤ 0.05 and if a log2 fold-change of 0.5 or greater was observed. Data are deposited in the NCBI Gene Expression Omnibus (GEO) database under Accession no. GSE200320). Heatmaps (all significantly altered genes with FDR<0.05) were build using with GENE-E v3.0.215. In order to do pathway enrichment analysis, Ingenuity Pathways Analysis (IPA) (43) was used focusing on differentially expressed genes with an FDR of < 0.05. As determined from the DESeq2 differential gene expression analysis above. In order to predict transcription factors that regulate Ces family genes, iRegulon V1.3 (44) was run on the cytoscape 3.8.0 platform (45).

### Serine Hydrolase Probe MS Experiments

For serine hydrolase probe MS experiments (13) primary enterocytes were isolated from 5 mice to prepare a single sample (N=1) and pooled cells together to extract protein by sonication in PBS. Protein concentrations were measured by micro BCA or Bradford assay. 300-400 µg of protein at a concentration of 1 mg/ml was incubated with either Fluorophosphonate-biotin probe (final concentration of 5µM) or equivalent amount of DMSO (as negative control) for 60 minutes at 37ºC. Excess probe was removed and protein precipitated with chloroform/methanol extraction by adding 2 volumes of methanol, 0.5 volume of chloroform and 1 volume of H_2_O and subsequently vortexed and centrifuged at 14,000 rpm for 5 minutes. The top layer was discarded and the protein layer collected from tube bottom. 2 volumes of methanol were added to the protein and stored it in -80ºC overnight. The following day, the protein pellet was centrifuged, excess methanol removed and the protein pellet air dried for 15 minutes. The protein pellet was resuspended in freshly prepared 500µl 6M urea in 25 mM ammonium bicarbonate followed by the addition of 2.5 µl 1mM DTT and incubation at 65ºC for 15 minutes. After cooling, 20µl 0.5M iodoacetamide was added to the protein which was then incubated at room temperature for 30 minutes to alkylate free cysteines. 70 µl of 10% SDS was added and heated for 5 minutes at 65ºC. Samples were diluted with 3 ml PBS and incubated with 50 µl streptavidin-agarose beads at room temperature for 2-3 hours on a shaker. Beads were precipitated by centrifuging at 2500Xg for 2 minutes, washed, and resuspended in 250 µl 25 mM ammonium bicarbonate. 1 µg trypsin was added per sample and incubated overnight on a shaker at 37ºC. Samples were then centrifuged and the supernatant containing peptides was collected followed by peptide desalting through C18 columns. Peptides were quantified and 200ng of sample loaded onto instrument for LC-MS analysis.

### Mass Spectrometry Analysis

A nanoElute was attached in line to a timsTOF Pro equipped with a CaptiveSpray Source (Bruker, Hamburg, Germany). Chromatography was conducted at 40°C through a 25cm reversed-phase C18 column (PepSep) at a constant flowrate of 0.5 μL/min. Mobile phase A was 98/2/0.1% Water/MeCN/Formic Acid (v/v/v) and phase B was MeCN with 0.1% Formic Acid (v/v). During a 108 min method, peptides were separated by a 3-step linear gradient (5% to 30% B over 90 min, 30% to 35% B over 10 min, 35% to 95% B over 4 min) followed by a 4 min isocratic flush at 95% for 4 min before washing and a return to low organic conditions. Experiments were run as data-dependent acquisitions with ion mobility activated in PASEF mode. MS and MS/MS spectra were collected with m/z 100 to 1700 and ions with z = +1 were excluded.

Raw data files were searched using PEAKS Online Xpro 1.6 (Bioinformatics Solutions Inc., Waterloo, Ontario, Canada). The precursor mass error tolerance and fragment mass error tolerance were set to 20 ppm and 0.03 respectively. The trypsin digest mode was set to semi-specific and missed cleavages was set to 2. mouse Swiss-Prot reviewed (canonical) database (downloaded from UniProt) and the common repository of adventitious proteins (cRAP, downloaded from The Global Proteome Machine Organization) totaling 20,487 entries were used. Carbamidomethylation was selected as a fixed modification. Oxidation (M) was selected as a variable modification.

Experiments were performed in biological triplicate. Resulting combined datasets were subjected to the following filtration criteria:

1. Database Search (−10log(p-value) ≥ 20, 1% peptide and protein FDR).
2. cross-reference with a serine hydrolase proteome dataset.
3. Generate ratio of Probe/No Probe. Require ≥2 Unique Peptides and ≥3 peptides total with probe treatment
4. Proteins determined to be probe enriched were 3-fold more detected in Probe-treated sample compared to No Probe (ratio of ≥3)

Data is available via the UCSD Mass Spectrometry Interactive Virtual Environment, a full member of the Proteome Exchange consortium, under the dataset number (Accession no. MSV000089304).

### Protein isolation and western blot

Primary enterocytes were centrifuged in PBS and the cell pellet was incubated with protein lysis buffer (20mM Tris-HCl pH8.0, 137mM NaCl, 1% Nonidet P-40 (NP-40) and 2mM EDTA) overnight in -80°C before protein isolation was carried out by repeated cycles of freezing and thawing. To isolate protein from tissue samples, a TissueLyser (Qiagen) was used to homogenize the tissue samples in lysis buffer. Lysates were then centrifuged at 13,000×g for 10 minutes at 4°C to pellet debris and supernatants were stored in -80°C for future use. Protein concentration was measured by Bradford assay, followed by western blotting using standard procedure. 10-20 µg protein samples in SDS-PAGE were resolved in 7.5% -10% gels (Bio-Rad) and transblotted onto polyvinylidene fluoride membranes (Millipore). Membranes were blocked with 5% BSA-PBST for 1 hour and then incubated with primary antibody (listed in supplementary table 1) overnight at 4°C. Membranes were then washed in 0.15% PBST 3-5 times at 5 minutes per wash before incubation with HRP-conjugated secondary antibodies for 1 hour. Membranes were washed 3-5 times in 0.15% TBST. Immunoreactive bands were generated using an Immobilon Western chemiluminescence HRP-conjugated substrate (Amersham) and developed either on a film (Kodak) or imaged in a ChemiDoc. For figure 2D, Li-Cor secondary antibodies were used to generate bands and imaged in Li-Cor Odyssey. Membranes were subsequently deprobed using Restore western blot stripping buffer (Thermo scientific) and re-probed using other primary antibodies

### Immunofluorescence staining

Mice jejunal tissues were fixed in 4% Z-fix overnight followed by cryopreservation in 15% and then 30% sucrose in PBS. Tissues were then embedded in OCT medium and cryo-sectioned (30 µm) on frost-free slides using a cryo-stat. Coverslips/ slides were then washed with PBST (0.5%Tween or Triton-X-100) and incubated in blocking buffer (PBST, 1% bovine serum albumin and 5% donkey serum) for 1 hour. Tissue sections were incubated overnight with the primary antibody against Epcam (primary antibodies and their dilutions are listed in supplementary table 1) in blocking buffer at 4°C. On the following day, tissue sections were washed with PBST 3 times and then incubated with the secondary antibodies (donkey anti rat Alexafluor in 1:100 dilution) for 1 hour, washed, stained with Bodipy 488 (2mg/ml) for 30 minutes followed by mounting in Vectashield (H-1400) DAPI. For staining for active hydrolases, the fixed tissues on slides were preincubated for 20 minutes with assay buffer (50 mM Tris-HCl, pH 7.4; 1 mM EDTA; 100 mM NaCl; 5 mM MgCl2 and 0.1% (w/v) BSA) followed by incubation for 60 min with TAMRA-FP in the assay buffer (0.5 μM final concentration)(12). Slides were then washed 3 times in 0.1M phosphate buffer before mounting with DAPI. Images were captured in the confocal microscope Zeiss LSM 780-FLIM and processed in Image J.

### Microsome isolation

A microsome isolation kit (ThermoFisher Scientific) was used for separating cytosol and microsome from jejunal tissues following the manufacturer’s protocol. Jejunal tissues from mice fed with radiolabeled oleic acid were resuspended in 200 µl of olive oil. 50 mg tissue was homogenized in homogenizing buffer, incubated on ice for 1 hour and centrifuged at 10,000g for 10 minutes to clear debris. The supernatant was centrifuged at 20,000g for 20 minutes in the pellet containing the microsomes, washed and resuspended in resuspension buffer with the supernatant representing the cytosolic fraction. Radioactivity was measured in each of these fractions using liquid scintillation counting.

### Human small intestine samples

Human intestinal epithelial cells from small intestinal resection tissue samples from inflammatory bowel disease patients (46) were provided by Dr. Rieder, Cleveland Clinic Foundation, with appropriate IRB approval. Intestinal tissue pieces were lysed using RIPA buffer followed by protein quantification by micro-BCA assay and 30 µg protein samples were loaded in 10% polyacrylamide gels for western blotting.

### Acute fat challenge

For acute fat challenge experiments, mice were fasted for 4 hours and then subjected to an oral gavage of olive oil (200 µl). Mice were euthanized 2 hours after the oil bolus and intestinal tissue pieces were collected for further experiments. Mice were treated IP with lipoprotein lipase inhibitor Tyloxapol (0.5 mg/g body weight of mice) 1 hour before the oral gavage (47). WT mice were treated with FATP2 blocker Grassofermata (Cayman chemicals, catalog no. 26202) by IP injection at a dose of 300mg/kg 2 hours prior to oral gavage (48). WT mice were injected IP with either β5 blocking antibody (5 mg/kg) or isotype control antibody 2 hours before acute fat challenge. Blood was drawn from mouse tail veins before and 2 hours after oil gavage.

### ^3^H oleic acid gavage

4 hour-fasted mice were subjected to oral gavage with olive oil containing 5 µCi ^3^H-labeled oleic acid. Mice were treated IP with lipoprotein lipase inhibitor Tyloxapol (0.5 mg/g body weight of mice) 1 hour before oral gavage. Prior to and 30, 60 and 120 minutes after olive oil/^3^H oleic acid administration, blood was collected from the tail vein. Mice were then euthanized and intestinal tissue pieces were procured and then freeze-dried in a tissue lyophilizer.

### Chronic high-fat feeding

8-week-old male mice were placed on high-fat diet (60%kcal% fat, Research Diet Inc, catalog no. d12492) for 3 weeks after which they were fasted for 12 hours before evaluation of the serum and intestinal TG content.

### Serum and intestinal TG measurement

Serum and intestinal TG content was measured using TG measurement kit (Cayman Chemical) following manufacturers’ protocol. Blood was collected from mouse tail veins and serum isolated by centrifuging blood samples for 15 minutes at 2000×g. 5 µl serum were used to measure TG content. For measuring TG content in the intestinal samples, approximately 50 mg tissue were homogenized in lysis buffer (supplied with the kit) and used 10 µl of the tissue lysate to measure TG content. The total TG content were normalized to the weight of the tissue.

### Ex-vivo overexpression of HNF4γ

An adenoviral expression vector containing mouse *Hnf4γ* gene (pAV[EXP]-EGFP CMV>mHnf4g (NM_013920.2), referred to here as AD-*Hnf4γ*-GFP) was obtained from Vector Biolabs. The intestinal lumen was flushed with PBS and pieces of small intestine from *Hnf4γ* KO mice were incubated with either blank adenoviral vector or *Hnf4γ*-OV for 24 hours in DMEM. Intestinal pieces were then washed with PBS and lysed for protein extraction and western blot.

### Luciferase assay

The 500-bp region of the putative promoter/enhancer regions of multiple Ces1 and Ces2 genes (corresponding to the ChIP-Seq peak, accession no. XXXX)(15) were subcloned into a luciferase vector with a minimal promoter (PGL4) and dual luciferase activity assay was performed in control and HNF4γ-overexpressed HEK293 cells in the presence and absence of oleic acid. HEK293 cells were plated on 96-well plates and infected with Hnf4γ-expressing adenovirus for 6 hours in serum-free media and then cells were incubated with complete media overnight. On the following day, Hnf4γ-adenovirus infected cells were transfected with 0.5 μg of reporter plasmid carrying the firefly luciferase gene under the control of Ces gene promoters containing the HNF4γ binding sequences in PGL-4 vector, and 0.5 μg of reference plasmid pRL-TK carrying the Renilla luciferase gene under the control of the simian virus 40 enhancer and promoter (Promega). Lipofectamine 3000 reagent (Invitrogen) was used as a transfection reagent following the manufacturer’s protocol. After 24 hours, cells were treated with oleic acid (0.6 µM final concentration) for another 12 hours. Cells were lysed in 200 μl of passive lysis buffer (Promega). Firefly luciferase and Renilla luciferase activities were measured using Dual-Luciferase reporter assay system (Promega) on a 96-well plate on a plate reader. Relative activity was defined as the ratio of firefly luciferase activity to Renilla luciferase activity.

### HNF4γ protein stability assay

HEK293 cells were infected with Hnf4γ-adenovirus for 6 hours in serum-free media and then the cells were incubated with complete media overnight. On the following day, the cells were first treated with cyclohexamide (100 µg/ml) followed by treatment with oleic acid in fat-free BSA or BSA alone. Cells were then lysed 6, 12 and 24 hours after treatment. Protein samples were used to detect HNF4γ protein level by western blotting.

## Acknowledgement

This work was supported by awards from the NIH (HL136377-01 and DK110098) to K.A. R.D. was supported by the Larry L. Hillblom Foundation Fellowship Research Grant (2019-D-004-FEL). The sequencing was carried out at the UC Davis Genome Center DNA Technologies and Expression Analysis Core, supported by NIH Shared Instrumentation Grant 1S10OD010786-01. The graphical abstract was made using Biorender.com. We would like to thank S. Layer for ongoing inspiration.

## Figure legends

**Supplementary Figure 1:**
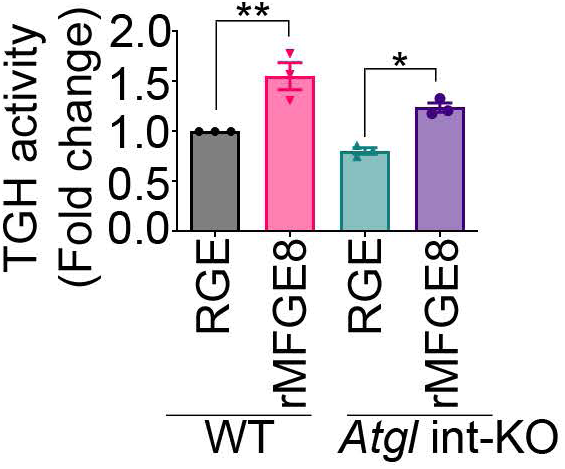
MFGE8 regulates enterocyte TG hydrolase activity independent of ATGL. TG hydrolase activity in WT and *Atgl* int-KO primary enterocytes after treatment with rMFGE8 or RGE control for 1 hour. N=3 independent experiments. Data expressed as Mean ± S.E.M. \**P* < 0.05, \*\**P* < 0.01. Data were analyzed by one-way ANOVA followed by Bonferroni’s posttest.

**Supplementary Figure 2:**
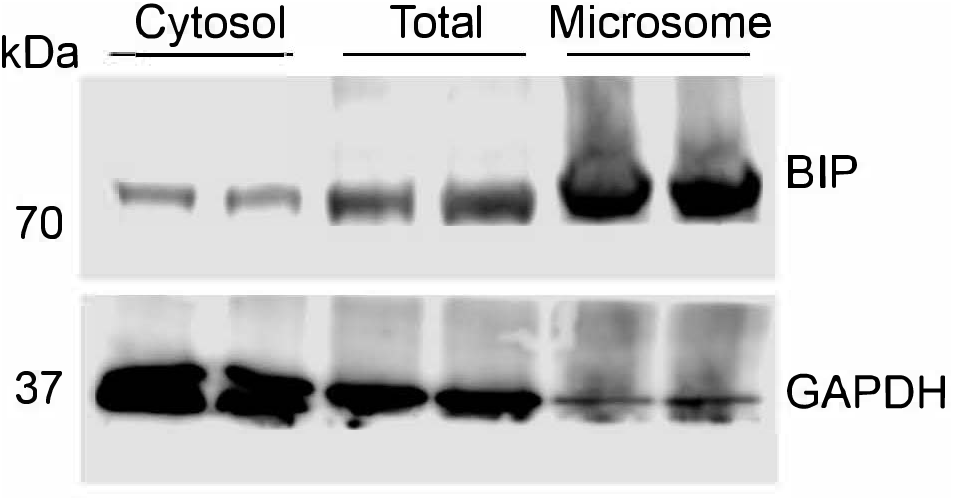
Validation of cytosolic and microsomal fractionation from total intestinal lysate. Western blot showing relative enrichment of microsomal (BIP) and cytosolic marker (GAPDH) proteins in respective subcellular fractions.

**Supplementary Figure 3:**
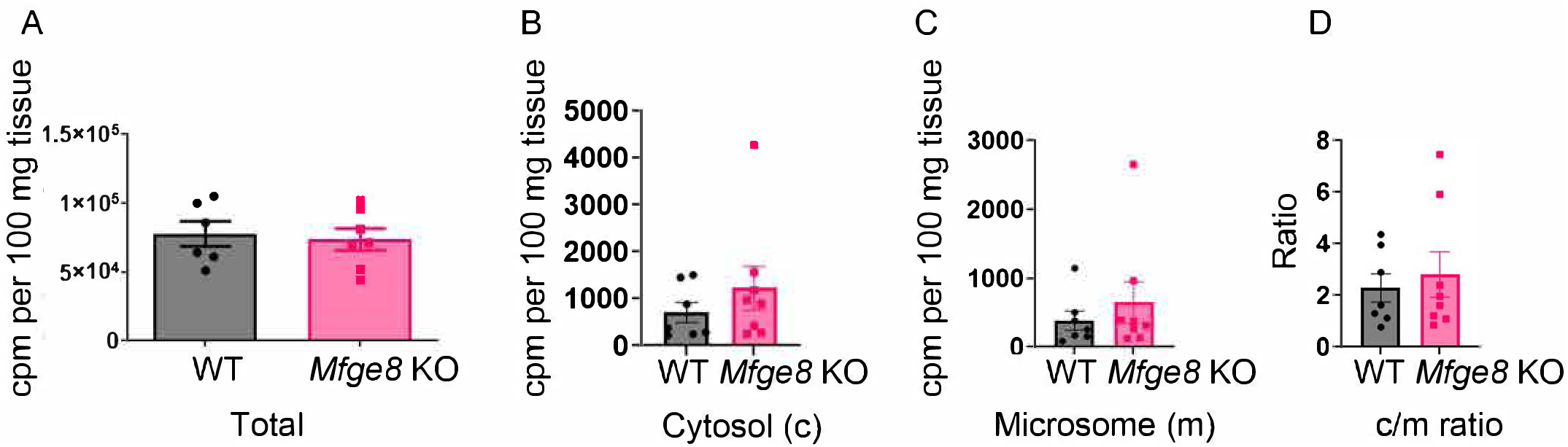
MFGE8 does not impact hydrolysis of cLDs derived from the basolateral surface. **(A-C)** ^3^H signal in the total (A), cytosolic fraction (B), microsomal fraction (C), and the ratio of cytosolic to microsomal radioactive signal (D) in the small intestines from WT and *Mfge8* KO mice 2 hours after i.p injection of ^3^H-labeled oleic acid. N=6-8 mice in each group. Data represents merged data from 2 independent experiments. Data expressed as Mean ± S.E.M.

## Graphical abstract

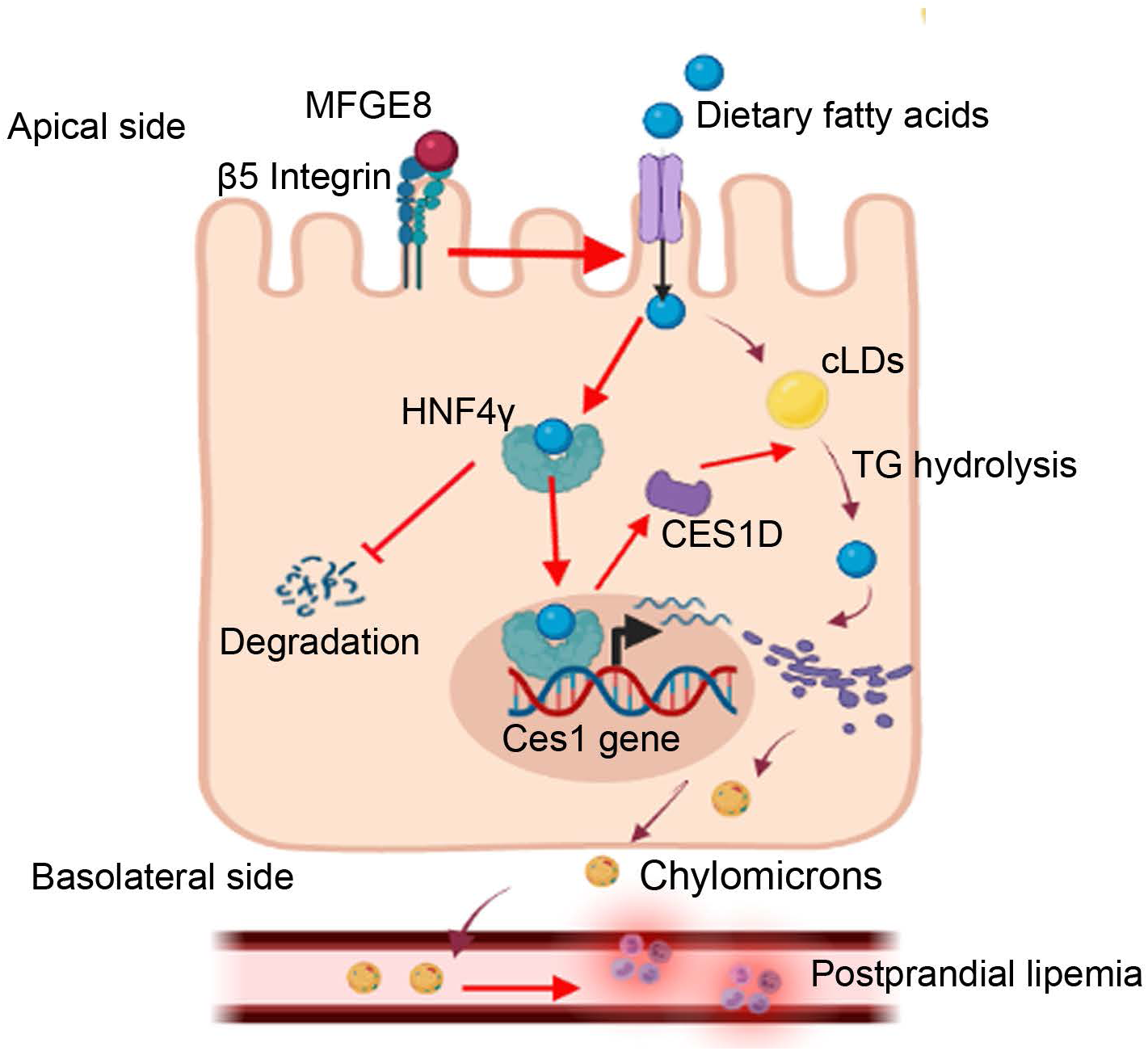

**Supplementary Table 1:**
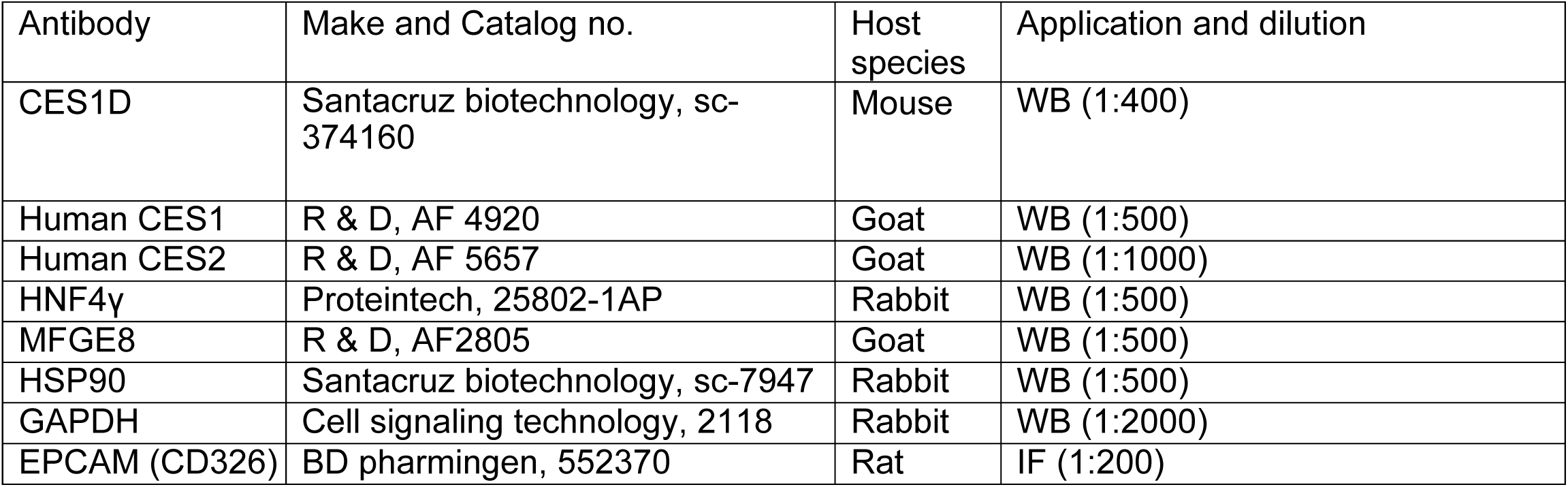
List of primary antibodies.

